# Nanobodies equipped with HaloTag variants enable rapid and straightforward one-step immunofluorescence lifetime multiplexing

**DOI:** 10.64898/2026.03.02.709025

**Authors:** László Albert, Samrat Basak, Henrike Körner, Nazar Oleksiievets, Nikolaos Mougios, Elena R. Cotroneo, Michelle S. Frei, Jörg Enderlein, Johannes Broichhagen, Nadja A. Simeth, Roman Tsukanov, Felipe Opazo

## Abstract

Multiplexed fluorescence imaging is limited by spectral overlap, whereas fluorescence lifetime provides an orthogonal encoding dimension. We replace genetically fused HaloTags with recombinant nanobody-HaloTag constructs applied as immunofluorescence reagents and determine a lifetime palette. Integrating lifetime and spectral encoding enables up to eight targets in a single acquisition and supports multiplexed imaging in cells and tissue using a simple OneStep labeling strategy compatible with fluorescent proteins, STED nanoscopy, and any standard laboratory antibody.

## Main Text

Fluorescence lifetime (FL) depends on the direct chemical environment of fluorophores and is sensitive to factors such as pH and nearby amino acids, which introduces complexity to FL Imaging Microscopy (FLIM). However, Frei and colleagues^1^ exploited this environmental sensitivity by mutating the wild-type HaloTag (HT7) to generate variants (HT9, HT10, and HT11) that alter the local chemical environment of the fluorophore, thereby producing distinct and separable FL signatures. This elegant strategy enables multiplexed imaging of up to three proteins of interest (POIs) in living cells by genetically encoding distinct HT fusions and decoding them via FLIM after labeling with a single fluorescent HaloTag ligand (HTL). More broadly, FL multiplexing has been demonstrated previously using various dyes and advanced computational or phasor-based analysis pipelines^2–6^. However, in most of these implementations, multiplexing relied on specific organelle-targeting dyes (e.g., mitochondria and nucleus), which limits flexibility: while multiple cellular compartments could be resolved, specific POIs could not be freely addressed without extensive chemical redesign or genetic manipulation. As a result, FL multiplexing was technically feasible but not readily adaptable to protein-centric biological questions. Here, we overcome these limitations by fusing all HT variants to nanobodies (Nbs) and extending FL multiplexing to any nanobody- or antibody-addressable POIs without genetic manipulation of the target proteins. Furthermore, we exploit the principle that spectrally similar or even identical fluorophores can reproducibly exhibit distinct FLs, enabling simple and scalable multiplexing (Figure 1a).

**Figure 1.**
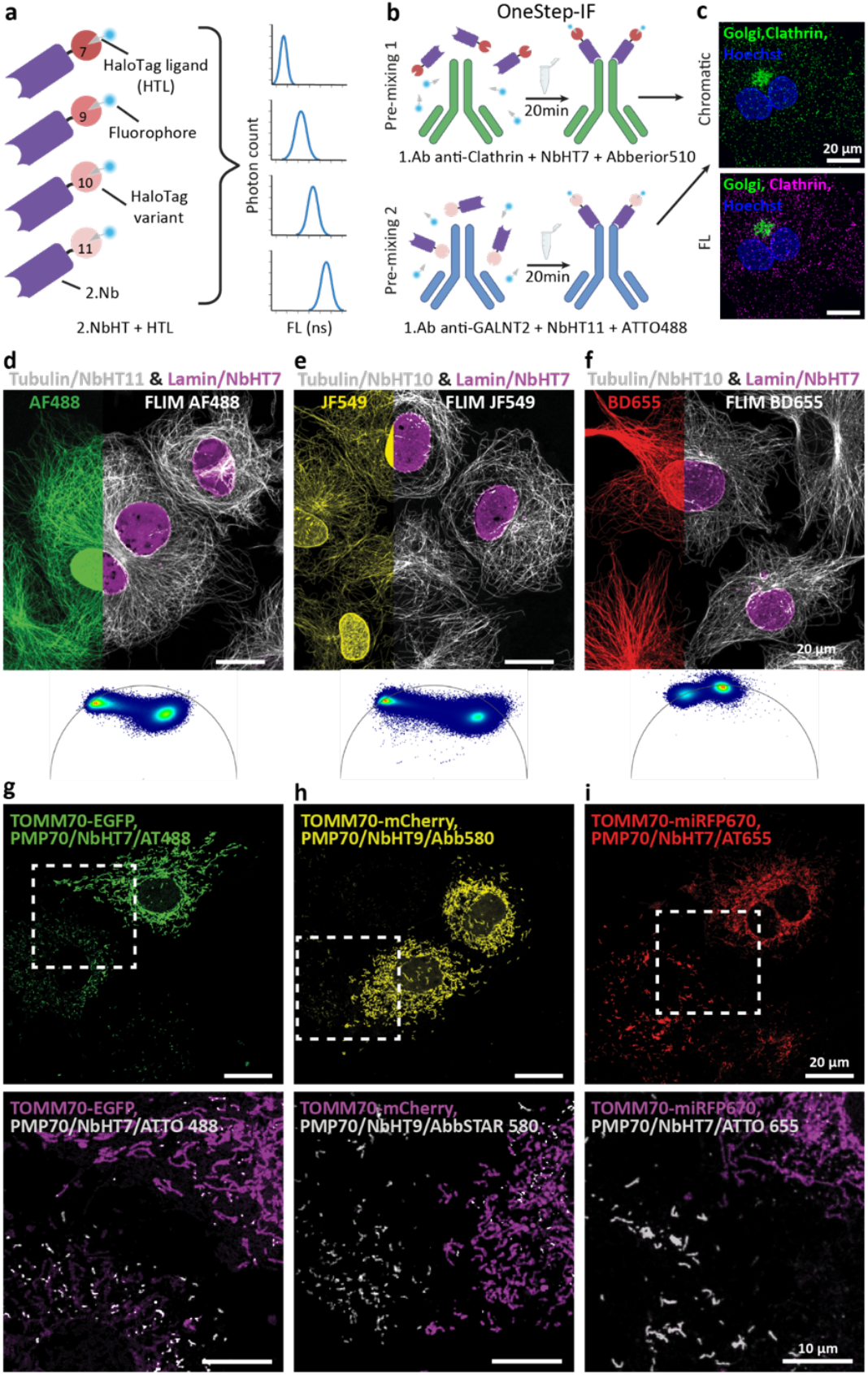
NanoFLex enables one-step, species-independent immunofluorescence through HaloTag–nanobody FLIM multiplexing. (**a**) Conceptual schematic illustrating that identical fluorophores conjugated to HTLs exhibit distinct FL when bound to different HT variants, enabling separation within a single spectral channel. (**b**) OneStep-IF workflow using 1.Abs pre-mixed with 2.NbHT allows simultaneous staining of multiple targets in a single incubation step. (**c**) Example of FL target separation: two rabbit 1.Abs against clathrin and a Golgi protein GALNT2 are labeled with spectrally overlapping ATTO488 and Abberior510 HTL conjugates via distinct HT variants, which are chromatically indistinguishable but readily separated by FLIM. (**d-f**) Benchmarking of HTL combinations (See Supp. Fig. 1) identifies fluorophore HT pairs suitable for robust FL separation. Representative phasor plots illustrate well-separated lifetime populations of a single fluorophore. (**g-i**) NanoFLex, in combination with genetically encoded fluorescent proteins, demonstrates that our approach is not limited to IF or to organic fluorophores.

Nanobodies fused to HaloTag variants for FL multiplexing (NanoFLex) provide several advantages, especially by allowing the flexible detection of multiple POIs within a chromatic channel for any target that can be immunoassayed. It would be ideal to fuse the HT variant to primary Nbs (1.Nbs); however, the palette of 1.Nbs working in immunofluorescence (IF) of cells or tissue is relatively small. Therefore, we considered strengthening this approach by using secondary Nbs (2.Nbs^7–9^), which allow for the pre-mixing of primary antibodies (1.Abs) with 2.Nbs in a OneStep-IF (Figure 1b). This strategy allows the use of multiple 1.Abs from the same species on the same sample^7,10^. Thus, two 1.Abs from the same species can be used in parallel, as each is recognized by a 2.Nb fused to a different HT variant (2.NbHT). While the fluorophores on the HTL can be the same or spectrally overlap, remaining indistinguishable by chromatic detection, the HT variants encode distinct FLs, enabling robust separation in the lifetime domain. (Fig. 1c) Our strategy showcases how simple and straightforward it is to perform multiple OneStep-IF with the preferred antibody of any laboratory in a species-independent manner. To expand the concept, we first benchmarked 26 fluorophores conjugated to HTL, creating a database of FL for all four HT variants (Supp. Fig. 1). Using this information, we selected the best combination to perform the initial test to separate FL when imaged with a commercial FL confocal laser-scanning microscope capable, employing routinely used laser lines for excitation (i.e., 488, 561, and 640 nm). To demonstrate that this strategy can work with 1.Nb as well as with 2.Nbs, we used COS-7 cells transiently transfected with LaminA fused to a fluorescently deficient mCherry (Y71L-mCherry). The cells were stained with an anti-mCherry 1.Nb fused to HT7 in combination with pre-formed complexes of anti-alpha-Tubulin 1.Abs and 2.NbsHT10 or 2.NbHT11 (Fig. 1d-f).

This simple evaluation showed that the concept works not only with 1.Nbs and 2.Nbs, but also allowed precise localization of two targets per excitation wavelength, converting a routine three-channel image into a 6-plex with a single staining step.

The concept also works in Tau-STED mode (Supp. Fig. 2); however, FL with STED is complex, since the stimulated emission depletion beam alters the fluorophores’ lifetimes, generating a FL gradient from the center of the depletion donut (zero intensity) to its maximal intensity^11^.

Fluorescent proteins (FPs) are widely used as labeling methods for a range of specimens; therefore, we evaluated their compatibility with NanoFLex. Hence, we transiently transfected COS-7 cells to produce the transmembrane domain of TOMM70 fused to mEGFP, mCherry, or miRFP670^12^ to decorate the cytosolic surface of mitochondria. We then performed OneStep-IF of PMP70 on peroxisomes using fluorescently labeled HTLs whose emission spectra overlapped with those of the corresponding FPs. Despite the chromatic overlap, the labeled targets were successfully separated in the FL domain (Fig. 1g-i). Most FPs have relatively well-defined short and narrow FL distributions (mEGFP, 2.07±0.05 ns; mCherry, 1.35±0.09 ns; miRFP670, 1.61±0.02 ns; mean±SD), which makes their combination with NanoFLex a straightforward procedure.

Although three FLs can be resolved within a single spectral channel^2,13–15^, and our system is technically capable of performing such separation, in practice, under our typical imaging conditions, FL distributions partially overlap. This results in photons emitted by different HaloTag variants contributing to the same pixel during acquisition. When diverse emitters are present within a single pixel, their combined decay reduces the separability of lifetime populations, particularly in densely labeled regions. This increased overlap compromises robust target discrimination and can lead to misclassification or exclusion of ambiguous pixels. To minimize this risk and ensure robust subcellular interpretation, we restricted the analysis to two well-separated FL populations per spectral channel (≥0.5 ns separation), thereby improving the reliability of photon classification and spatial assignment in our system. With this approach, we use a pre-mix of 1.Abs with 2.NbsHT or 1.NbHT. The staining was performed in approximately 1h, and with using the species independent OneStep-IF we revealed 8-targes, including vimentin (intermediate filaments), α-Tubulin (microtubules), TOMM20 (mitochondria), PMP70 (peroxisomes), NUP50 (nuclear pore complex), GALNT2 (Golgi apparatus), LaminA-Y71L-mCherry (nuclear envelope), and clathrin (coated pits; Fig. 2a-e). We further validated the approach by performing six-target detection, together with nuclear staining (DAPI), in brain slices (Fig. 2f–i). To further enhance the flexibility of our approach, we combined HaloTag-fused 2.Nbs with directly fluorophore-conjugated 2.Nbs, allowing implementation even when a desired fluorophore is not available as HTL. Two targets per spectral channel were resolved by pairing HaloTag-loaded 2.Nbs (AbbLive510, Sulfo549, or SiR HTLs) with directly dye-conjugated 2.Nbs (AbbSTAR Green, azDye 568, or Alexa Fluor 647). Despite strong spectral overlap between dye pairs, distinct FL enabled robust separation in tissue.

**Figure 2.**
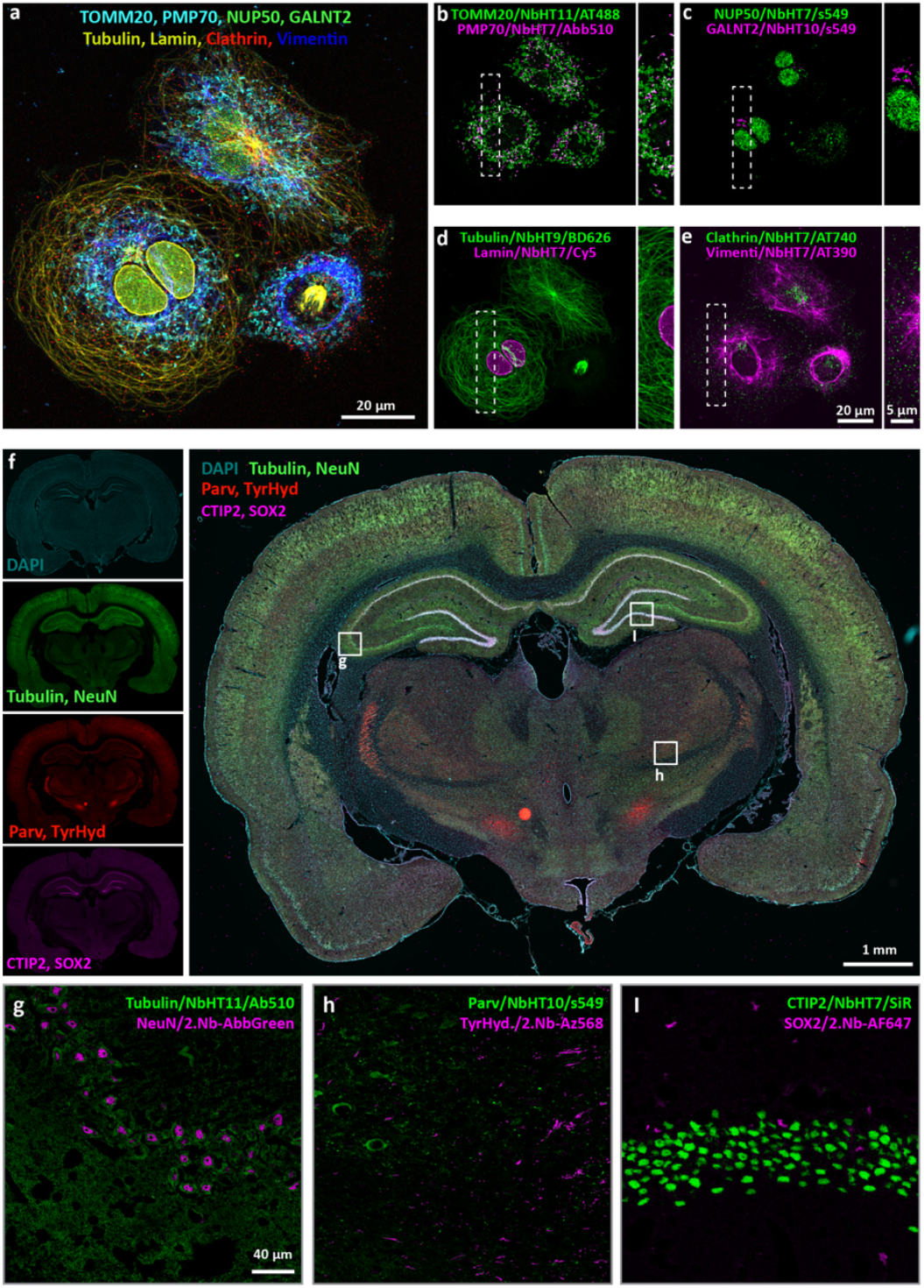
Multiplex NanoFLex in cells and tissue using a single staining step. (**a-e**), Eight subcellular structures labeled in COS-7 cells using OneStep-IF and resolved by FLIM. Two targets are separated per spectral channel using distinct HaloTag variants conjugated to chromatically overlapping fluorophores. HTL-ATTO390 and ATTO740 are chromatically separated. Targets include Vimentin, TOMM20, PMP70, NUP50, GALNT2, LaminA-Y71L-mCherry, and Clathrin. (**f–i**) Six-target NanoFLex plus DAPI in rat brain cryosections demonstrates compatibility with fixed tissue and directly conjugated 2.Nbs. All images were acquired on a confocal FLIM platform and separated using phasor-based lifetime analysis.

In summary, we introduced NanoFLex, a universal FL multiplexing strategy that combines HaloTag variants with nanobody-based immunolabeling to achieve high-plex detection in a single staining step. The approach is fully compatible with standard antibodies, species-independent multi-labelling, fixed cells, tissue sections, fluorescent proteins, and STED super-resolution microscopy, without requiring genetic manipulation of the sample. By extending from the spectral to the lifetime domain, this method transforms routine immunofluorescence into scalable, chromatically efficient, and experimentally accessible high-content imaging. We anticipate that 2.Nbs-HaloTag FL multiplexing will be readily adopted as a drop-in extension of existing antibody workflows across cell biology.

## Material and Methods

### Cell culture

COS-7 cells were cultured in a humidified incubator at 37 °C with 5% CO2 in Petri dishes, using complete Dulbecco’s Modified Eagle Medium (DMEM, ThermoFisher Scientific) supplemented with 10% fetal bovine serum (FBS, ThermoFisher), 4 mM L-glutamine, and 1% pen/strep (ThermoFisher) antibiotics. For immunostainings, cells were seeded in 12-well plates on 18 mm poly-L-lysine-coated coverslips and incubated for 16-18 hours in the same incubator.

### Transient Transfection

Approximately 16 hours before fixation, cells were transfected in 12-well plates using Lipofectamine 2000 (Thermo Scientific) transfection kits according to the manufacturer’s recommendations. For each well, 300 ng of plasmid was diluted to 50 μl using OPTIMEM medium (Gibco) in one Eppendorf tube, while 2 μl of Lipofectamine 2000 reagent was added to another Eppendorf tube containing 48 μl of OPTI-MEM. After thorough mixing, the contents of the two tubes were combined and incubated for 20 minutes and then added to each well and plates were placed back to the incubator (37°C, and 5% CO2) for 16-20 hours.

### Rat Brain Cryosections

Lewis rats on a LEW/Crl background (Rattus norvegicus) were bred at the animal facility of the University Medical Center Göttingen (Germany). The rats were kept under standardized conditions on a 12-h light–dark cycle. They were provided with food and drink ad libitum. All experiments were performed in accordance with local animal welfare regulations. A female Lewis rat was intracardially perfused with saline, followed by a fixative containing 4% paraformaldehyde (PFA). Brain tissue was postfixed in 4% PFA for 24 hours and immersed in 30%. Brain tissue was embedded in OCT medium, and 18 µm thick cryosections were cut for staining.

### Synthesis of fluorescently-labeled HaloTag ligands

For ATTO390-HTL, ATTO-488-HTL, ATTO740-HTL, ATTO655-HTL, Cy5.5-HTL, and Cy7-HTL, commercially available NHS-ester derivatives of fluorophores (1 mg each) were transferred into a 1.5 mL Eppendorf tube using MeCN and diluted to a total volume of 1 mL. Then, a solution of HTL-NH_2_ (2-(2-((6-chlorohexyl)oxy)ethoxy)ethan-1-amine; 1.1 molar equivalent with respect to the selected NHS-fluorophore in a minimum amount of MeCN) was added to the solution. The reaction mixture was shielded from light and placed on a shaker at ambient temperature for 18h, while the process of the reaction was followed through HPLC-MS analysis (Agilent 1260 Infinity II combined with an InfinityLab LC/MSD iQ mass detector (ESI) using an InfinityLab Poroshell 120 EC-C18 column at 40 °C eluted with MeCN/H_2_O + 0.1% formic acid). After completion, the reaction mixture was directly subjected to HPLC purification using a JASCO system (PU-4086 pumps, a UV-4075 detector; MonoQTM HR 5/5 column with a flow rate of 3 mL/min, 30 to 70% MeCN (+0.1% TFA) in MilliQ (+0.1% 15 TFA) gradient over 25 minutes). The identity of the product was confirmed using HPLC-MS (ESI) and the fraction containing the product was lyophilized (Christ Alpha 2-4 LD plus lyophilizer). ATTO390-HTL: calc. for C_30_H_46_ClN_2_O_5_^+^ [MH^+^] 549.3, found 349.4; ATTO-488-HTL: calc. for C_35_H_44_ClN_4_O_11_S_2_^+^ [MH^+^], 795.2 found 795.3; ATTO740-HTL: structure not disclosed, calc. 673.3, found 673.4, ATTO655-HTL: calc. for C_37_H_54_ClN_4_O_7_S^+^ [MH^+^] 733.3, found 733.4; Cy5.5-HTL: calc. for C_50_H_59_ClN_3_O_15_S_4_^3-^ [M^3-^] 368.1, found 368.2; Cy7-HTL: calc. for C_47_H_65_ClN_3_O_9_S_2_^+^ [MH^+^] 914.4, found 914.2.

For the generation of the Sulfo608-HTL, AbberriorLIVE510-HTL, AbberiorSTAR580-HTL, AbberiorLIVE590-HTL, ATTO590-HTL, CF594-HTL, ATTO655-HTL, Cy5-HTL, most chemical reagents and anhydrous solvents for synthesis were purchased from commercial suppliers (Sigma-Aldrich, Roth) and were used without further purification unless otherwise stated. Alexa Fluor488-HTL has been purchased from Promega (#G1001), HaloTag Ligands linked to Sulfo549/646^16^, ATTO700^17^, iFluor647^18^, and TMR/SiR^19^ have been described before. UPLC-UV/Vis was performed on an Agilent 1260 Infinity II LC System equipped with an Agilent SB-C18 column (1.8 µm, 2.1 × 50 mm). Buffer A: 0.1% FA in H_2_O Buffer B: 0.1% FA in acetonitrile. The typical gradient started with 10% or 30% B for 1.0 min, then ramped to 95% B over 5 min. Finally, it was maintained at 95% B for 1.0 min with 0.6 mL/min flow. Preparative or semi-preparative HPLC was performed on an Agilent 1260 Infinity II LC System equipped with columns as follows: preparative column – Reprospher 100 C18 columns (10 μm: 50 × 30 mm at 20 mL/min flow rate; semi-preparative column – 5 μm: 250 × 10 mm at 4 mL/min flow rate. Eluents A (0.1% TFA in H_2_O) and B (0.1% TFA in MeCN) were applied as a linear gradient. Peak detection was performed at the maximal absorbance wavelength. A carboxylic acid-containing dye (30 nmol) was dissolved in DMSO (150 µL), and DIPEA (120 nmol) and TSTU (33 nmol, from a 10 mg/mL stock solution in DMSO) were added sequentially, and the solution was incubated at room temperature for 15 minutes. HTL-NH_2_ (33 nmol) was added, and the reaction was allowed to stand for 1 h before quenching with JB Special solution^20^ (MeCN:dH2O:HOAc = 25:25:1; 100 µL). The reaction was then subjected to RP-HPLC, collecting at the respective maximal wavelength, and the resulting mixture was lyophilised. Compounds HRMS Sulfo608-HTL: HRMS (ESI): calc. for C_46_H_59_ClN_5_O_13_S_2_ [M]^+^: 988.3, found: 988.3. AbberiorLIVE510-HTL HRMS (ESI): calc. for C_35_H_35_ClF_8_N_3_O_6_ [M]^+^: 780.2, found: 780.3. AbberiorSTAR580-HTL Structure not disclosed. AbberiorLIVE590-HTL HRMS (ESI): calc. for C_36_H_45_ClN_3_O_5_ [M]^+^: 634.3, found: 634.3. ATTO590-HTL HRMS (ESI): calc. for C_47_H_59_ClN_3_O_6_ [M]^+^: 796.4, found: 796.5. CF594-HTL Structure not disclosed. ATTO655-HTL HRMS (ESI): calc. for C_37_H_54_ClN_4_O_7_S [M]^+^: 733.3, found: 733.4.Cy5-HTL HRMS (ESI): calc. for C_42_H_59_ClN_3_O_3_ [M]^+^: 688.4, found: 688.5.

### Fluorescence lifetime determination

Fluorescence lifetime data were analyzed using Leica LAS X FLIM software. FLIM images were acquired using time-correlated single-photon counting (TCSPC) with sufficient photon counts to ensure smooth fluorescence decay curves and well-defined phasor distributions. Excitation and acquisition parameters were adjusted to maintain sufficient photon detection while avoiding the detection of multiple photons per laser pulse, thereby preventing pile-up effects. Lifetime analysis was performed using predominantly default software settings without binning, applying either the wavelet or preview filters with threshold values set between 1 and 3. Lifetime-based channel separation for multiplexed imaging was achieved by resolving distinct lifetime populations in the phasor plot. For Atto390 and DAPI, excitation was performed using a continuous-wave (CW) diode laser, which does not allow fluorescence lifetime detection on this system.

### Immunofluorescence of cells

For fixation, COS-7 cells were first treated with pre-warmed Extraction Buffer (EB, 138 mM KCl, 10 mM MES, 3 mM MgCl2, 2 mM EGTA, 320 mM sucrose, pH 6.8) supplemented with 0.1% Saponin for 25s. The solution was then removed and immediately replaced with pre-warmed (37°C) Fixation Buffer (4% PFA, 0.1% Glutaraldehyde in EB) and incubated for 15 minutes at room temperature. After removing the fixatives, the cells were rinsed once with PBS (137 mM NaCl, 2.7 mM KCl, 10 mM Na2HPO4, 2 mM KH2PO4, pH 7.4), and free aldehyde groups were quenched with 0.1 mM Glycine in PBS for 15 minutes. Fixed and quenched cells were then blocked and permeabilized with 3% bovine serum albumin (BSA) and 0.1% Triton X-100 (Sigma Aldrich) in PBS for 30 minutes at room temperature. Before each immunolabeling, all primary antibodies (1.Ab) were separately pre-mixed with their corresponding Halo-tagged 2.Nbs (2.NbHT, custom-made by NanoTag Biotechnologies GmbH) and fluorescent HaloTag ligands were incubated in a 1:3:4 molar ratio in 15 µl PBS for 20 minutes at room temperature or overnight at 4°C. The list of all antibodies and nanobodies used is available in the Supp. Tables 1 and 2. The resulting staining complexes were pooled and adjusted to the desired volume with blocking buffer (PBS and 3% BSA), then incubated with the cells for 1h at room temperature with gentle agitation. Samples were washed and rinsed three times with PBS for 5 minutes each, briefly dipped in distilled water, mounted in Mowiol (12 ml 0.2 M Tris buffer pH 8.5, 6 ml ddH_2_O, 0,6 g glycerol, 2.4 g Mowiol, Carl Roth, cat. no. 0713.2)., and dried for 0.5h at room temperature.

### Immunofluorescence of tissue

Tissue sections were removed from the freezer and incubated at 37 °C for 10 minutes to warm up and dry the samples. The embedding medium was removed by one wash with PBS, and the glass around the tissue was carefully wiped dry. In the dry area, the sections were encircled using a liquid blocker pen and left to dry for 1 minute, while avoiding the tissue itself to dry. The tissue sections were then placed in a humidified chamber and blocked with 8% BSA supplemented with 0.5% Triton X-100 and 0.2% Tween-20 for 1 hour. In the meantime, primary antibodies, Halo-tagged 2.Nbs, and fluorescently labeled HaloTag ligands were premixed separately in 15-20 µl PBS in separate Eppendorf tubes at a molar ratio of 1:3:4, respectively, and incubated at room temperature (RT) for at least 30 minutes. Antibodies and nanobodies are found in the Supp. Tables 1 and 2. After incubation, the staining complexes were pooled together, and the volume was increased to 250 µl to achieve final concentrations of 5% BSA and 0.05% Triton X-100 in PBS. The blocking solution was then removed, and the samples were incubated with 200–220 µl of the staining solution overnight at 4 °C in a humidified chamber. The following day, slides were rinsed once with PBS and incubated with DAPI in PBS for 5 minutes. The specimens were then washed three times for 5 minutes each in PBS. The glass around the tissue was wiped clean to remove residual blocking lipids, and the tissue was mounted in 15–20 µl Mowiol and dried at RT. Fluorescent overview images of tissue sections were taken with a VS200 slide scanner (Olympus) and 20x air objective (UPLXAPO20X). Subsequently, images were taken with confocal microscopy and FLIM-based channel separation.

### Microscopy and image analysis

Confocal FLIM imaging was performed on a STELLARIS 8 PP STED FALCON microscope (Leica Microsystems, Wetzlar, Germany) equipped with an HC PL APO 100×/1.4 OIL STED W objective. Excitation in the visible range was provided by a supercontinuum white-light pulsed laser (80 MHz), whereas DAPI was excited using a 405 nm diode laser. Excitation (and corresponding detection) wavelengths for the GFP, Cy3, Cy5, and NIR channels were 497 nm (510–560 nm emission), 560 nm (571–607 nm emission), 649 nm (660–730 nm emission), and 743 nm (753–830 nm emission), respectively. When changing fluorescent dyes, the excitation wavelength and detection window can be adjusted analogously to classical confocal microscopy without affecting fluorescence lifetime imaging measurements. Emission in the DAPI, visible, and near-infrared ranges was detected using HyD S, HyD X, and HyD R hybrid detectors, respectively, all operated in photon-counting mode. After locating the region of interest, FLIM mode was activated. Images were acquired with a pixel size of 110 nm and a pixel dwell time of 10 µs. Line accumulation (1–6) and laser power were adjusted individually to optimize signal-to-background ratio for each set of fluorophores. Following imaging parameters optimization, a z-stack of 5–8 optical sections was acquired with a 0.4 µm step size. Fluorescence lifetime images were analyzed and represented in the lifetime phasor space. Separation of multiplexed targets was achieved by selecting the corresponding lifetime populations directly in the phasor plot. For Atto390 and DAPI, excitation was performed using a diode laser operating in CW mode. Custom confocal FLIM measurements (Supp. Fig. 1) were performed using a home-built confocal setup, as described previously^15^ (Supp. Fig. 3). Briefly, excitation was provided by a white-light (WL) laser (Koheras SuperK Power) operated at a fixed repetition rate of 40 MHz. An acousto-optic tunable filter (AOTF) (Koheras SpectraK Dual) enabled flexible selection of the excitation wavelength. The laser beam was coupled into a single-mode fiber (SMF) (PMC-460Si-3.0-NA012 3APC-150-P, Schafter+Kirchhoff) using a fiber coupler (60SMS-1-4-RGBV-11-47, Schafter+Kirchhoff) and collimated after fiber output using a 10× air objective (UPlanSApo 10×, 0.40 NA, Olympus). A quad-band dichroic mirror (ZT405/488/561/640rpc, Chroma) directed the excitation light to the sample and separated excitation and emission light. The excitation beam was scanned using a fast laser scanning system (FLIMbee, PicoQuant GmbH), allowing imaging of extended regions of interest while maintaining a fixed focal plane by focusing the laser beam on the back plane of the objective (UApo N 100×, 1.49 NA oil, Olympus). Lateral positioning and axial focusing were controlled manually with a XY stage (Olympus) and a z-piezo stage (Nano-ZL100, Mad City Labs), respectively. Fluorescence emission was collected by the same objective, focused through a 100 µm pinhole (P100S, Thorlabs) using a 180 mm achromatic lens (AC508-180-AB, Thorlabs), and subsequently collimated by a 200 mm lens (AC508-200-A, Thorlabs). Long-pass filters (488 LP or 647 LP Edge Basic, Semrock) suppressed residual excitation light, while band-pass filters (BrightLine HC 692/40, 609/54, or FF 536/40; Semrock) further reduced scattered light detection. The emission light was then focused onto a single-photon avalanche diode detector (SPCM-AQRH, Excelitas) using an achromatic lens (AC254-030-A-ML, Thorlabs). Photon arrival times were timed using a TCSPC module (HydraHarp 400, PicoQuant). Data acquisition and synchronization of the TCSPC and scanning systems were controlled using SymPhoTime 64 software (PicoQuant). For CLSM-based FL-SMLM measurements, 80 × 80 µm^2^ regions of interest (ROIs) were scanned. Each image consisted of four repeated scans with a virtual pixel size of 100 nm and a pixel dwell time of 20 µs. For confocal FLIM data analysis, the custom Matlab-based software kit TrackNTrace (lifetime edition) was used^21^. The software is freely available on GitHub via the link: https://github.com/scstein/TrackNTrace. For lifetime value extraction, the first 0.1 ns after the maximum of the TCSPC histograms were discarded, and the TCSPC tail was fitted with a mono-exponential decay using a maximum likelihood estimator (MLE), as described previously. Then lifetime average values and distribution widths have been extracted from lifetime histograms generated from lifetime values in each pixel.

## Authors contribution

L.A., S.B., H.K., N.O., N.M., E.C., N.A.S., and R.T., performed the experiment, generated brain slices, determined the fluo-lifetimes, and synthesized the fluorescent HTL. M.S.F., J.B., provided material. J.E., provided equipment and advised. R.T., and F.O., supervised the project. F.O., conceived and led the project. L.A., R.T., and F.O., wrote the manuscript. All authors revised and helped edit the manuscript.

## Acknowledgments

We thank Jannik Hentze for great technical support, Kai Johnsson for initial discussion and support, Zhixing Chen for providing dyes conjugated to HTL, Steffen Frey and Hansjörg Götzke for support and producing these HaloTag fusion at NanoTag Biotechnologies GmbH. The authors thank Ingo Gregor for help with lifetime data analysis as well as target separation procedure. The authors thank Jan Christoph Thiele for continuous support with the confocal FLIM data analysis and modifying the Matlab-based routines. We also thank Julia Najderek, Dawid Pęksyk, and Jędrzej Przybył (all FMP) for assistance with fluorescence HaloTag ligands. F.O. was supported by Deutsche Forschungsgemeinschaft (DFG) through the SFB 1286 (project Z04). H.K. was funded by the Deutsche Forschungsgemeinschaft (DFG) – SFB TRR 274/2 2024 – 408885537, SFB 1690 – 528760423 and from the European Commission under the European Union’s Horizon 2020 research and innovation programme, grant agreement No 101021345.

## Notes

### Competing Interest Statement

F.O. is a shareholder of NanoTag Biotechnologies GmbH. The other authors declare no competing interest

