## Supplementary material for "Nanobodies equipped with HaloTag variants enable rapid and straightforward one-step immunofluorescence lifetime multiplexing": Supp Fig.

**Supplementary Table 1.** List of all antibodies.

| Target | Host | Isotype | Clonality | Company | Cat. No. | RRID |
|---|---|---|---|---|---|---|
| Tubulin | mouse | IgG1 | mono | Synaptic Systems | 302211 | AB_887862 |
| Vimentin | mouse | IgG1 | mono | Santa Cruz | sc-6260 | AB_628437 |
| Tomm20 | mouse | IgG1 | mono | Sigma Aldrich | WH0009804M1 | AB_1843992 |
| PMP70 | rabbit | IgG | mono | Abcam | ab85550 | AB_10672335 |
| Clathrin | rabbit | IgG | poly | Abcam | ab21679 | AB_2083165 |
| GALNT2 | rabbit | IgG | poly | Sigma Aldrich | HPA011222 | AB_1849446 |
| NUP50 | rabbit | IgG | mono | Abcam | ab137092 | AB_2921286 |
| Parvalbumin | mouse | IgG1 | mono | Synaptic Systems | 195011 | AB_2619883 |
| NeuN | rabbit | IgG | mono | Synaptic Systems | 266108 | AB_3662033 |
| CTIP2 | rabbit | IgG | mono | Abcam | ab240636 | AB_10730034 |
| vGlut1 | rabbit | IgG | poly | Synaptic Systems | 135303 | AB_887875 |
| Tyrosine hydrox. | mouse | IgG2 | mono | Synaptic Systems | 213111 | AB_2636902 |
| SOX2 | rabbit | IgG | poly | Synaptic Systems | 347003 | AB_2620099 |

**Supplementary Table 2.** List of all nanobodies.

| Target | Host | Clonality | Fusion/Conj. | Company | Cat. No. | RRID |
|---|---|---|---|---|---|---|
| Mouse IgG1 | alpaca | mono | Atto643 | NanoTag Biotechnologies | N2002-At643-S | AB_3076022 |
| Mouse IgG2 | alpaca | mono | azDye658 | NanoTag Biotechnologies | N2702-AF568-S | AB_3076052 |
| Rabbit IgG | alpaca | mono | AlexaFluor647 | NanoTag Biotechnologies | N2402-AF647-S | AB_3076036 |
| Rabbit IgG | alpaca | mono | AbberiorSTAR Green | NanoTag Biotechnologies | N2402-AbGREEN-S | AB_3697322 |
| RFP/mCherry | alpaca | mono | HT7 | NanoTag Biotechnologies | custom made | - |
| Mouse IgG1 | alpaca | mono | HT7 | NanoTag Biotechnologies | N2441 | AB_3668679 |
| Mouse IgG1 | alpaca | mono | HT9 | NanoTag Biotechnologies | custom made | - |
| Mouse IgG1 | alpaca | mono | HT10 | NanoTag Biotechnologies | custom made | - |
| Mouse IgG1 | alpaca | mono | HT11 | NanoTag Biotechnologies | custom made | - |
| Mouse IgG2 | alpaca | mono | HT7 | NanoTag Biotechnologies | N2741 | AB_3668678 |
| Rabbit IgG | alpaca | mono | HT7 | NanoTag Biotechnologies | N2041 | AB_3668677 |
| Rabbit IgG | alpaca | mono | HT9 | NanoTag Biotechnologies | custom made | - |
| Rabbit IgG | alpaca | mono | HT10 | NanoTag Biotechnologies | custom made | - |
| Rabbit IgG | alpaca | mono | HT11 | NanoTag Biotechnologies | custom made | - |

a

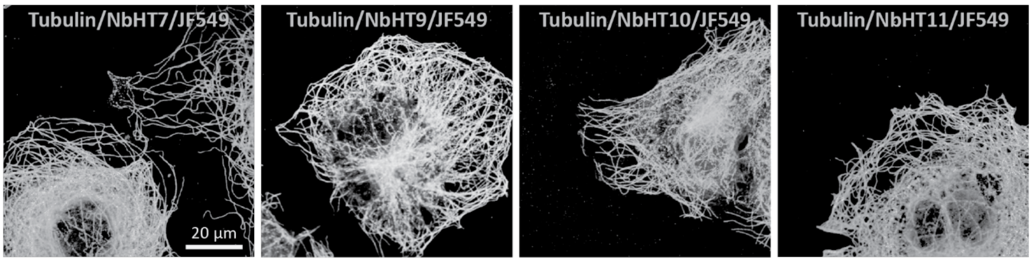


b

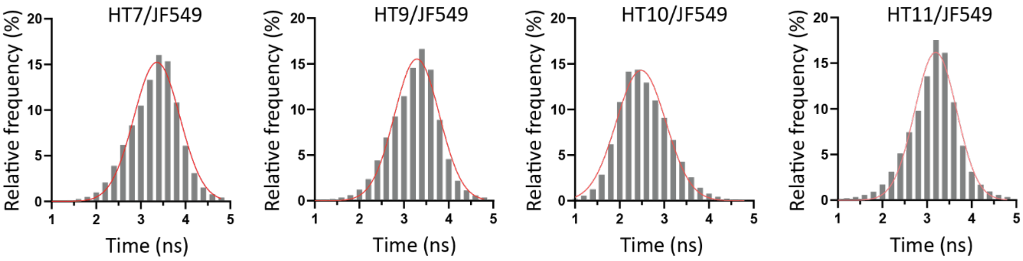


c

| HaloTag ligand | HaloTag-7 | HaloTag-9 | HaloTag-10 | HaloTag-11 |
|---|---|---|---|---|
| AlexaFluor 488 | 3.10±0.72 | 3.19±0.58 | 2.50±0.75 | 1.36±0.46 |
| ATTO 488 | 2.85±0.94 | 2.96±0.77 | 2.20±0.72 | 1.88±0.36 |
| abberiorLIVE 510 | 2.98±0.82 | 2.62±1.30 | 1.93±0.62 | 0.90±0.25 |
| JaneliaFluor 549 | 3.36±0.52 | 3.29±0.50 | 2.48±0.55 | 3.19±0.47 |
| Sulfo 549 | 3.16±0.54 | 3.23±0.73 | 1.77±0.37 | 2.77±0.45 |
| TMR | 2.85±0.40 | 2.81±0.39 | 2.18±0.35 | 2.64±0.38 |
| BD555 | 3.07±0.54 | 3.04±0.74 | 2.20±0.58 | 2.81±0.53 |
| MaP555 | 3.06±0.61 | 2.92±0.61 | 2.24±0.92 | 2.75±0.46 |
| abberiorSTAR 580 | 3.04±1.09 | 3.44±0.91 | 3.35±0.77 | 3.36±0.93 |
| abberiorSTAR 590 | 3.48±0.64 | 3.35±0.75 | 1.83±0.80 | 2.99±0.61 |
| ATTO 590 | 3.32±0.66 | 3.44±0.75 | 3.16±0.58 | 3.29±0.73 |
| CF594 | 2.90±0.44 | 2.88±0.59 | 2.43±0.40 | 2.24±0.40 |
| Sulfo608 | 3.02±0.57 | 3.03±0.62 | 1.84±0.28 | 2.01±0.36 |
| MaP618 | 3.25±0.61 | 3.53±0.71 | 2.02±0.33 | 2.36±0.50 |
| BD626 | 2.77±0.91 | 2.77±1.08 | 2.24±0.91 | 2.70±0.96 |
| Sulfo646 | 3.25±0.56 | 3.26±0.60 | 1.52±0.22 | 2.38±0.55 |
| iFluor647 | 1.94±0.27 | 1.90±0.48 | 1.70±0.43 | 1.81±0.34 |
| SiR | 3.16±1,11 | 3.40±0,60 | 2.52±0.56 | 2.88±0.67 |
| Cy5 | 1.60±0.17 | 1.67±0.22 | 1.74±0.28 | 1.65±0.19 |
| ATTO 655 | 2.40±0.35 | 2.18±0.39 | 2.11±0.30 | 1.50±0.33 |
| BD655 | 3.11±0.44 | 2.99±0.45 | 2.12±0.25 | 2.65±0.36 |
| Cy5.5 | 1.73±0.49 | 1.71±0.23 | 1.60±0.37 | 1.63±0.22 |
| ATTO 700 | 2.11±0.34 | 2.19±0.35 | 1.68±0.36 | 1.73±0.30 |
| SiR 700 | 2.66±0.28 | 3.24±0.56 | 2.56±0.38 | 2.47±0.33 |
| Cy7* | 0.92±0.01 | 0.92±0.00 | 0.94±0.01 | 0.94±0.01 |
| ATTO 740* | 0.89±0.01 | 0.94±0.01 | 0.83±0.04 | 0.73±0.00 |

**Supplementary Figure 1.** Benchmarking 26 fluorophores conjugated to HTL. (a) OneStep-IF using anti-alpha-Tubulin 1.Ab combined with 2.NbHT7, HT9, HT10, or HT11 and in the example added HTL-JF549 on COS-7 cells. (b) FL distribution of full images and fit (red line) of the JF549 when combined with the different HaloTag variants, as shown in a. (c) Mean FL from the fits in ns and error are standard deviations for all tested fluorophores. All FLs were obtained using the setup described in the Methods section, except for Cy7* and Atto 740*, which were extracted from Leica STELLARIS software directly.

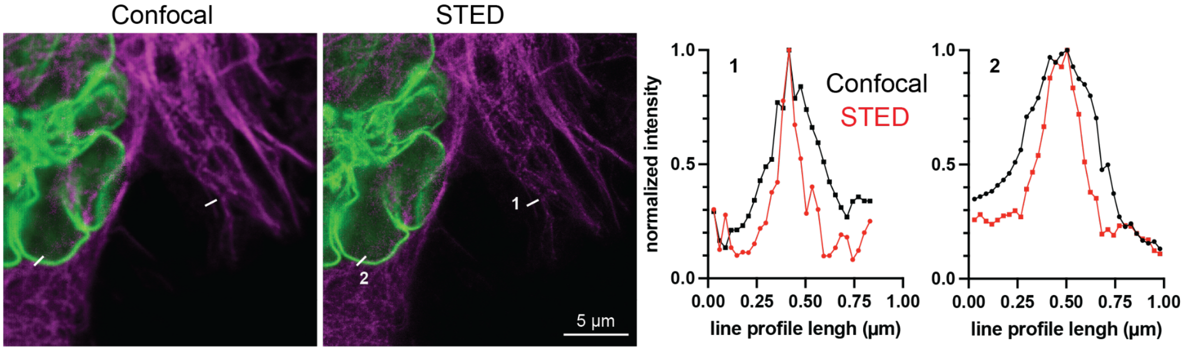


**Supplementary Figure 2.** Proof of concept that 2 targets on the same chromatic spectrum can be separated under Tau-STED. COS-7 cells transfected with LaminA-Y71L-mCherry (non-fluorescent mCherry, here in green) were stained with 1.Nb anti-RFP-HT7 and SiR-HTL (here displayed in green). Simultaneously, anti-alpha-Tubulin 1.Ab was combined with 2.Nb-Atto643 (here in Magenta). Line profile used to demonstrate the STED effect, separation of 2 targets imaged under the chromatic line.

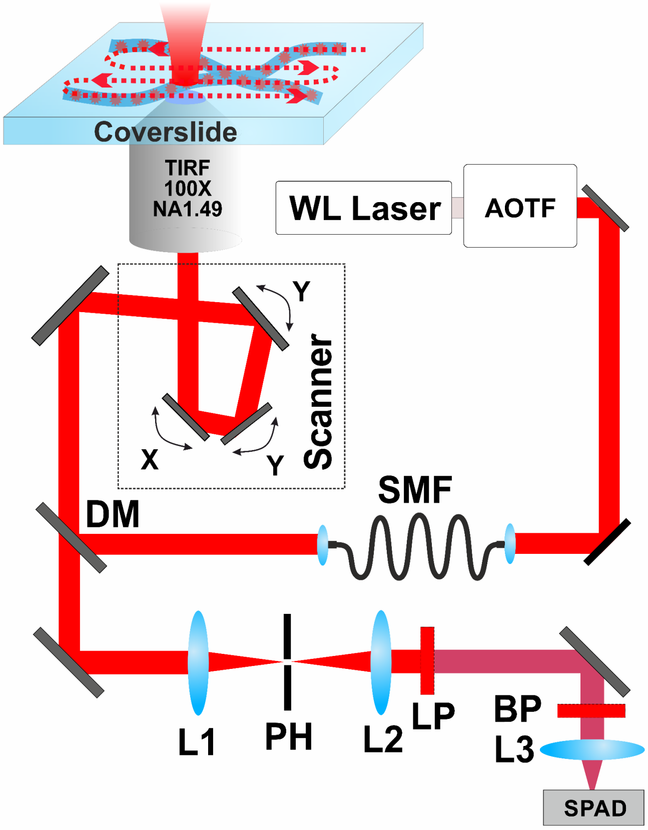


**Supplementary Figure 3.** Schematic of the custom-built time-resolved confocal laser-scanning microscope. Excitation is provided by a white-light (WL) laser and directed through a scanning confocal microscope to image microtubules immobilized on a glass coverslip. Fluorescence emission is collected confocally and detected by a single-photon avalanche diode for time-correlated single-photon counting.
